# Transcriptomic stress responses in *Vaccinium* spp. F1 hybrids: Implications for temperature-resilient cranberry breeding using a crop wild relative

**DOI:** 10.1101/2025.07.08.662648

**Authors:** Audrey Dickinson, Jackson Kubal, Nicholas Allsing, Emily Murray, Todd P. Michael, James J. Polashock

**Affiliations:** Department of Science and Conservation, San Diego Botanic Garden, 300 Quail Gardens Dr, Encinitas, CA, 92024-2707, USA; Plant Molecular and Cellular Biology, Salk Institute for Biological Sciences, La Jolla, California 92037, USA; Center for Marine Biotechnology and Biomedicine, University of California, San Diego, La Jolla, CA, 92093, USA; Department of Cell and Developmental Biology, School of Biological Sciences, University of California, San Diego, La Jolla, CA, 92093, USA; USDA, Agricultural Research Service, Genetic Improvement of Fruits and Vegetables Lab, 125A Lake Oswego Rd., Chatsworth, New Jersey 08019, USA

**Keywords:** abiotic stress, crop wild relatives (CWR), interspecific hybridization, temperature stress, transcriptomics, *Vaccinium macrocarpon* (American cranberry), *Vaccinium oxycoccos* (small cranberry)

## Abstract

Cranberry (*Vaccinium macrocarpon*) is an important North American fruit crop with vulnerability to temperature extremes and a relatively recent domestication history. Hybridization with a cold-adapted crop wild relative (CWR), *V. oxycoccos*, offers a strategy for improving temperature stress tolerance.
We conducted RNA sequencing (RNA-seq) on *V. macrocarpon* and F1 hybrids between *V. macrocarpon* and *V. oxycoccos* subjected to acute heat and cold stress, capturing early transcriptional responses up to 30 minutes (heat) and 95 minutes (cold) after treatment onset. We then evaluated differences in responses across genotypes and stress conditions.
Differential expression analysis and functional profiling revealed cold-induced differences in pathways related to photosynthesis, ribosomes, defense, and hormone signaling. No subgenome-specific functional specialization was observed. Two F1 hybrids exhibited suggestive cold resilience, with expression changes elevated at 60 minutes but declining by 95 minutes. Hybrids also displayed substantial regulatory variation under stress and transgressive downregulation of photosynthesis genes under ambient conditions.
These findings suggest that *V. oxycoccos* introgression could be utilized in breeding cold-tolerant cranberry cultivars. Variation observed between F1 hybrids reflects the diversity introduced through CWR germplasm and provides opportunities for selection. Conservation of *V. oxycoccos* and other CWRs remains critical for future crop improvement.

**Plain Language Summary:** Cranberries face increasing threats from temperature extremes. We explored crossbreeding cultivated cranberry (Vaccinium macrocarpon) and a cold-adapted wild relative (V. oxycoccos) to improve resilience to temperature stress. Hybrids showed different gene expression patterns from their parents, especially under cold stress. Our findings support the value of wild relatives in crop improvement and emphasize the importance of protecting wild plant diversity.

## Introduction

The United States is the world’s largest producer of cranberries (*Vaccinium macrocarpon* Aiton), a fruit crop of economic, cultural, and ecological significance. While cranberries have been commercially cultivated for over 200 years, indigenous people across North America have long harvested wild cranberries for food, medicine, and dye (Neto & Vinson, 2011; Turner & von Aderkas, 2012). Today, cranberries rank as the second most consumed berry in the United States after strawberries, primarily in processed forms such as juice and sweetened dried cranberries (Neto & Vinson, 2011; Bentley, 2018). Cranberries are also rich in bioactive compounds, including anthocyanins, proanthocyanidins, and flavonols. These have been associated with potential health benefits including urinary tract infection prevention and antioxidant activity (Neto & Vinson, 2011; Moro, 2024). The health effects of cranberry consumption and supplementation remain an area of active research, as reflected in dozens of ongoing and completed clinical trials (ClinicalTrials.gov, 2025). Concentrated in the Northeast, Wisconsin, and the Pacific Northwest of the United States (US), cranberry production supports a substantial agricultural sector responsible for $343.1 million in annual export volume in 2023 (Weber *et al*., 2024), encompassing a supply chain of growers, packers, and processors.

However, cranberry cultivation faces increasing challenges due to environmental stressors, particularly temperature extremes. Heat stress can directly damage fruit through scalding when ambient temperatures exceed 32°C (Croft, 1995; Oudemans & Mupambi, 2025), leading to reduced fruit quality and yield. Additionally, heat-induced scalding can also increase susceptibility to pathogens such as the complex of fungi that cause fruit rot, a major cause of economic loss in cranberry production (Oudemans & Mupambi, 2025). Cold stress introduces additional risk, as frost can severely damage sensitive developmental stages, particularly flower buds emerging from dormancy in spring and fruit ripening early in the fall, despite the general cold hardiness of dormant buds and mature fruit. Frost-damaged buds may fail to develop into fruit, while frost-damaged fruit may display reduced firmness and greater disease susceptibility (UMass Amherst Cranberry Station, 2019; DeMoranville, 1998).

Developing cranberry varieties with improved temperature resilience is therefore critical to maintaining sustainable production. Surveys of cranberry producers in the US and Canada highlight the importance of breeding for temperature stress tolerance. Among abiotic stress traits, heat tolerance was rated the highest priority by New Jersey producers, while tolerance to fall and spring frost was highest rated in Wisconsin and British Columbia (Gallardo *et al*., 2018). Further, the most desired agronomic traits overall (firmness, fruit size, anthocyanin content, and resistance to fruit rot) are all negatively impacted by temperature stress, either through direct physiological damage, developmental perturbation, or increased susceptibility to disease (DeMoranville 1998; Vorsa & Zalapa, 2019; Oudemans & Mupambi, 2025). These industry priorities emphasize the need for breeding efforts that target heat and cold stress tolerance in cranberry.

Introgression of wild germplasm is an established strategy for improving stress tolerance, disease resistance, and other agronomic traits in crops (Hajjar & Hodgkin, 2007; Dempewolf *et al*., 2017). Crop wild relatives (CWRs) harbor genetic diversity that can be used to introduce novel alleles that are absent in cultivated varieties, thereby enhancing resilience and expanding environmental adaptability of domesticated species. In addition to introducing specific adaptive genes, interspecific hybridization between a crop and its CWRs can result in heterosis, or hybrid vigor, which manifests as increased yield and stress tolerance due to elevated heterozygosity and novel epistatic interactions between divergent subgenomes (Hajjar & Hodgkin, 2007; Liu *et al*., 2024).

*V. oxycoccos,* a CWR of *V. macrocarpon*, is a promising source of genetic diversity for cranberry breeding. While morphologically distinct, the two species readily hybridize, producing somewhat fertile F1 progeny (Diaz-Garcia *et al*., 2021; Kawash *et al*., 2022). Both *V. macrocarpon* and certain populations of *V. oxycoccos* are diploid with highly syntenic genomes (Kawash *et al*., 2022), facilitating compatibility in crosses. The two species differ in their morphology and ecology: *V. oxycoccos* produces smaller berries and has a circumboreal distribution, whereas *V. macrocarpon* bears larger fruit and is adapted to northern-temperate regions. Although early allozyme-based studies suggested very low genetic diversity in wild *V. macrocarpon*, possibly reflecting a Pleistocene bottleneck (Bruederle *et al*., 1996), more recent SSR-based analyses indicate that both *V. oxycoccos* and wild *V. macrocarpon* exhibit substantial genetic variation, with *V. oxycoccos* exhibiting slightly greater diversity (Rodriguez-Bonilla *et al*., 2020). Both wild species thus retain potentially useful genetic diversity relative to the genetic homogeneity of cultivated varieties (Neyhart *et al*., 2022). *V. oxycoccos*’s adaptation to colder environments offers particular promise for improving cold tolerance in cultivated cranberry, while core stress genes active in both heat and cold response may enhance heat tolerance. However, the consequences of *V. macrocarpon* hybridization with *V. oxycoccos*, and the potential for introgression of stress-resilience traits to positively impact productivity, remain uncharacterized.

Cranberry breeding efforts have historically progressed slowly due to long generation times (7–8 years per cycle) and limited genetic diversity, with most cultivars being only a few generations removed from their wild progenitors (Vorsa & Zalapa, 2019). While past breeding programs have prioritized traits such as yield, fruit color, and disease resistance, there has been little explicit selection for environmental stress resilience, leaving cranberry production vulnerable to climate extremes. Advances in functional genomics offer new opportunities to accelerate breeding efforts by identifying candidate genes associated with stress responses. Recent genomic resources, including high-quality reference genomes for *V. macrocarpon* and *V. oxycoccos* (Diaz-Garcia *et al*., 2021; Kawash *et al*., 2022), provide a foundation for high-throughput molecular investigations. While past studies have focused on genetic mapping using molecular markers (Vorsa & Zalapa, 2019), comprehensive genome-wide or transcriptome-wide analyses remain limited, and, to our knowledge, no studies have yet characterized stress responses in cranberry using omics methods. Furthermore, *V. macrocarpon* x *oxycoccos* F1 hybrids have not been evaluated for emergent molecular characteristics, such as differentially regulated stress response pathways, which may arise from novel allelic combinations or regulatory perturbations within the hybrid genome.

In this study, we investigated the transcriptional responses of *V. macrocarpon* and interspecific F1 hybrids (*V. macrocarpon* × *V. oxycoccos*) to acute heat and cold stress. Given the contrasting native ranges of *V. macrocarpon* and *V. oxycoccos*, which are shaped largely by temperature, we hypothesized that these species harbor distinct regulatory networks for temperature perception and response. We propose that acute stress exposure can reveal transcriptional pathways involved in thermal adaptation. We present the first transcriptomic characterization of acute temperature stress responses in *V. macrocarpon*, alongside a comparative analysis of baseline and stress-induced gene expression in four F1 hybrids. Using differential expression and functional enrichment analyses, we assess the potential of *V. oxycoccos* germplasm to contribute adaptive regulatory variation that could be harnessed to improve temperature resilience in cranberry breeding.

## Materials and Methods

### Plant Material

Representatives of diploid populations of *V. oxycoccos* were collected by L. Bruederle in southeastern Alaska in 1996 (Mahy *et al*., 2000; L. Bruederle, unpublished notes). Representative accessions were collected as single vines. Four *V. oxycoccos* accessions (NJ96-20, N96-82, NJ96-95 and NJ96-127) were crossed with *V. macrocarpon* cultivars (Stevens, Ben Lear, Pilgrim) in 1998 to create the F1 hybrids. Pedigree information is provided in Table 1. Accession data for the *V. oxycoccos* plants, both those used as parents in F1 crosses and those used for RNA sequencing, is presented in Table S1. Each F1 hybrid used in this project consisted of randomly selected individuals from the progeny of a given cross. The cultivar Stevens was used as the *V. macrocarpon* representative. *Vaccinium oxycoccos* is a diminutive, relatively slow growing species, especially under New Jersey growing conditions. As such, *V. oxycoccos* plant material was only available for T0 (see temperature stress conditions below). Note that the F1 progeny should contain the organelles from the *V. macrocarpon* female parent, except for CNJ98-325-33 where *V.oxycoccos* (NJ96-20) was the female parent (Table 1). The F1 progeny were cloned, using cuttings, from the original seedlings. All plants were maintained in a greenhouse in Chatsworth, NJ USA) prior to the temperature stress experiments. Experiments were performed during the growing season in mid to late September, before cold acclimation begins.

**Table 1:**
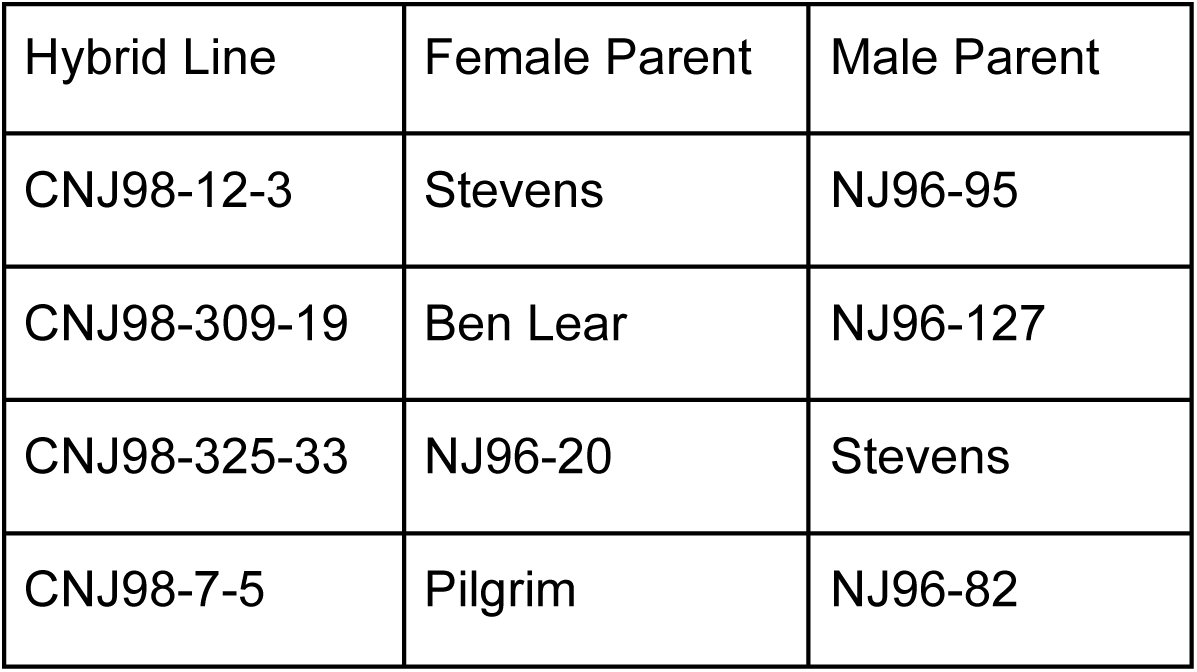
Pedigrees of Hybrid Lines. Each line is an interspecific hybrid between Vaccinium macrocarpon and V. oxycoccos, generated from a cross between the listed female and male parents. Cultivar names (e.g., ‘Stevens’, ‘Ben Lear’, ‘Pilgrim’) refer to V. macrocarpon cultivars, while NJ96-series identifiers represent V. oxycoccos accessions. Additional information on V. oxycoccos accessions is provided in Supplementary Table S1.

| Hybrid Line | Female Parent | Male Parent |
| --- | --- | --- |
| CNJ98-12-3 | Stevens | NJ96-95 |
| CNJ98-309-19 | Ben Lear | NJ96-127 |
| CNJ98-325-33 | NJ96-20 | Stevens |
| CNJ98-7-5 | Pilgrim | NJ96-82 |

### Temperature Stress Conditions

For acclimation prior to temperature stress, the F1 hybrids and the ‘Stevens’ *V. macrocarpon* representative were removed from the greenhouse and placed into incubators (Percival Scientific, Perry, IA USA). Identical incubators (Percival, 1-35VL) were used for both heat and cold treatment. Acclimation was at 21° C, with a 12 hour day/night cycle for three days prior to temperature stress experiments. There were three replicates per accession at each time point. For heat stress, incubators were set at 45° C with the plants already in the incubator. At time zero (T0), samples (whole leaves) were collected before the temperature began to go up. Leaves were snap frozen in liquid nitrogen and transferred to cold RNAlater ICE (ThermoFisher Scientific, Waltham, MA USA). Leaves were harvested at 15 m for T1 (actual incubator reading 39.7° C) and 30 m for T2 (actual incubator reading 47° C). Cold stress was done as noted above, but the incubators were set at 2° C. T0 samples were removed as soon as the temperature setting was lowered. T1 was harvested at 60 m (actual incubator reading 10.5° C). T2 was harvested at 95 m (actual incubator reading 3° C). We attempted to get as close to target temperature as possible, regardless of time in the incubator. For sample information, time points (T0, T1 and T2) are preceded by H or C to designate heat and cold treatment, respectively. See Figure 1 for a schematic of the experimental design.

**Figure 1:**
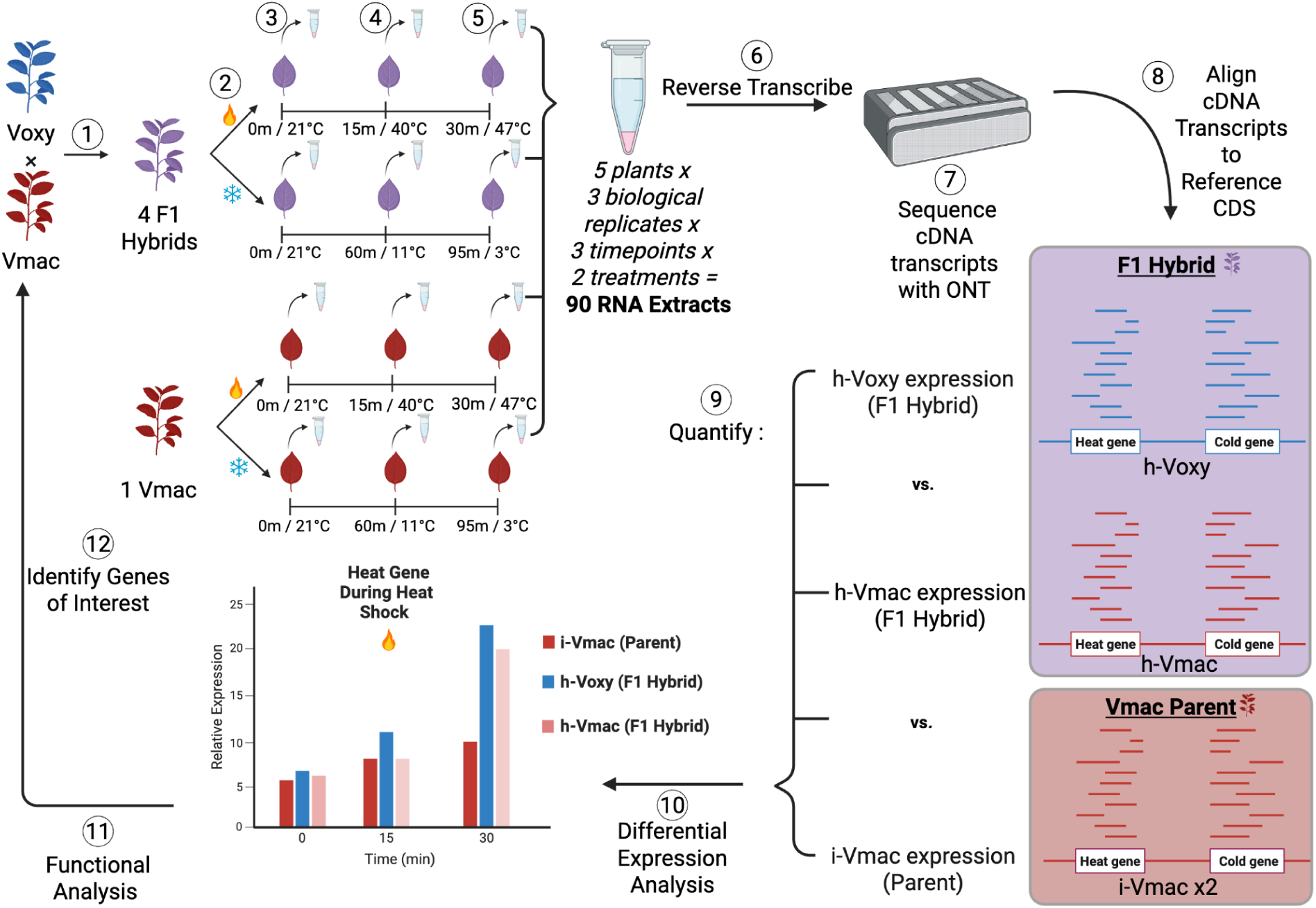
Temperature stress experimental design. Temperatures reached 47° C for heat treatment and 3° C for cold treatment. Differential expression analysis compared total gene expression in hybrids and V. macrocarpon by assigning genes to common KEGG ontology (KO) terms. A separate analysis compared the behavior of the V. macrocarpon subgenome in a pure (i-Vmac) versus hybrid (h-Vmac) context.

### RNA Isolation

Leaves were removed from the RNAlater ICE (4-5 leaves, about 50 mg) and processed to isolate total RNA. We used the CTAB procedure as noted in Daverdin *et al*., (2017) for total nucleic acid isolation. The total nucleic acid extractions were resuspended in 400 µl nuclease free water. RNA was isolated from the total nucleic acid by the addition of 100 µl 10M LiCl, followed by incubation on ice overnight. RNA was pelleted and resuspended in 180 µl nuclease free water. All samples were treated with DNAse (Optizyme, Fisherbrand #BP81071) according to the manufacturer’s instructions. Total RNA samples were then chloroform treated, precipitated with NaOAc and pelleted in a microcentrifuge. The pellet was washed with 70% ETOH, and finally resuspended in 25 µl nuclease free water for cDNA synthesis.

### Library Preparation and Sequencing

We employed full-length cDNA sequencing using Oxford Nanopore Technologies (ONT) to distinguish subgenome-specific expression in the F1 hybrids. The extended read lengths provided by ONT are expected to enhance mapping accuracy and resolution, thereby improving our ability to assign transcripts to their respective parental subgenomes. For each sample, 200 ng of RNA was used for full-length cDNA synthesis and PCR amplification using the ONT cDNA-PCR Sequencing Kit (SQK-PCB111.24) following the kit’s provided protocol. PCR amplification used barcoded primers and ran for 16 cycles. The resulting barcoded cDNA was pooled evenly into groups of nine samples per run. Adapters were ligated to the barcoded and pooled cDNA using the ONT Rapid Adapter Kit (EXP-RAA114) to ensure compatibility with V14 flow cell chemistry (EXP-AUX003) and loaded onto R10.4.1 flow cells for sequencing on the ONT PromethION platform. Sequencing was run for 72 hours. Sequencing runs were basecalled using the Dorado v5.0.0 super-high accuracy model with a quality threshold of Q9 and a 200 base pair cutoff. See Supplementary Data S1 for loading amounts and run statistics.

### Processing and mapping

Reference genomes for *V. macrocarpon* and *V. oxycoccos* were obtained from Kawash *et al*. (2022) and as posted in Genome Database for Vaccinium (GDV, vaccinium.org). To improve contiguity and enable chromosome-level comparisons, we scaffolded the *V. oxycoccos* genome using RagTag v2.1.0 (Alonge *et al*., 2022) with the *V. macrocarpon* assembly as a reference. Coding domain sequence (CDS) predictions for both genomes were generated using Helixer v0.3.2 (Stiehler *et al*., 2021). CDS completeness was evaluated using BUSCO v5.4.3 with the lineage dataset eudicots_odb10 (Manni *et al*, 2021). A hybrid reference CDS was created by concatenating CDSs from both genomes to facilitate mapping of reads from hybrid individuals. Reads were aligned to the corresponding CDS reference using minimap2 v2.28-r1209 (Li, 2018), with *V. oxycoccos* reads mapped to the *V. oxycoccos* CDS, *V. macrocarpon* reads mapped to the *V. macrocarpon* CDS, and hybrid reads mapped to the concatenated CDS (Supplementary Data S2). Only the best-matching primary alignment per read was retained. Transcript abundances were then quantified using Salmon v1.6.0 (Patro 2017). Full transcript quantification information is available in Supplementary Data S3.

### Functional annotation and ortholog detection

Coding domain sequences (CDSs) were annotated using eggNOG-mapper v2.0.1 and emapperdb v5.0.0 (Cantalapiedra *et al*., 2021), retrieving functional information including Gene Ontology (GO) terms, Kyoto Encyclopedia of Genes and Genomes (KEGG) pathway identifiers (Kanehisa *et al*., 2025), and orthologous group assignments. The *Arabidopsis thaliana* proteome was downloaded from Universal Protein Resources (UniProt) (The UniProt Consortium, 2025), and OrthoFinder v2.5.4 (Emms *et al*., 2019) was used to identify orthologous gene groups among *V. oxycoccos*, *V. macrocarpon*, and *Arabidopsis thaliana*, based on predicted protein sequences.

### Differential Expression and Functional Enrichment Analyses

Differential gene expression was analyzed using DESeq2 v1.44.0 in R v4.4.1 (Love *et al*., 2014). Samples were divided by species and experiment, with heat stress and cold stress experiments analyzed separately. The primary contrast compared the later stress condition to corresponding unstressed controls (HT2 vs. HT0 or CT2 vs. CT0). All comparisons used raw counts normalized via the median-of-ratios method, and genes were retained if they had ≥10 normalized counts in at least two samples. Genes with Benjamini-Hochberg FDR < 0.05 and absolute log2 fold change > 1 by a Wald test were considered differentially expressed. Full DESeq outputs were ranked by t-statistic, and gene set enrichment analysis (GSEA) was conducted on these gene sets via the clusterProfiler package v4.12.6 using the fgsea method with the number of permutations set to 100,000 (Subramanian *et al*., 2005; Yu *et al*., 2012; Korotkevich *et al*., 2021). Rank-based approaches such as GSEA enable the detection of functional enrichments in gene sets where significant gene-level responses are limited. GO term enrichment analysis was performed using the GSEA() function. KEGG pathway enrichment analysis was performed using the gseKEGG() function, and pathway information was accessed using the KEGGREST package (Tenenbaum & Maintainer, 2024). Pathways belonging to the “Human Diseases” KEGG classification were removed from enrichment results. Functional enrichments with a false discovery rate (FDR) < 0.05 were considered significant.

### Genomic Background Effects on *V. macrocarpon* Expression

We extracted expression data for *V. macrocarpon*-derived genes from both *V. macrocarpon* and hybrid samples to assess whether the *V. macrocarpon* subgenome responds to stress differently in a pure versus hybrid background. All genes were retained in *V. macrocarpon* samples, but raw counts corresponding to *V. oxycoccos* genes were discarded in hybrid samples before normalization to construct an *in silico* “hybrid-macrocarpon” transcriptome. Samples were split by experiment (HT0, HT1, and HT2 analyzed separately from CT0, CT1, and CT2), and DESeq2 results were extracted for the interaction term species*treatment, with a significant interaction effect defined as an adjusted p-value < 0.05. Gene lists were then ranked by t-statistic and analyzed by GSEA as described above.

### KEGG Pathway-Level Comparison

We collapsed transcript counts to KEGG orthology (KO) identifiers using eggNOG annotations to examine broad functional divergence between *V. macrocarpon* and hybrid transcriptomes. Counts were summed per KO term, and unannotated features were discarded. We then ran DESeq on this aggregated counts data, and results were extracted for the interaction term species*treatment to identify functional categories responding to stress differently between species. KEGG pathway enrichment analysis on the entire gene set was run using gseKEGG as described above, and microbial- or animal-specific pathways were removed after enrichment analysis prior to data visualization. Photosynthetic gene expression in each condition was quantified as the percentage of total normalized counts mapped to genes that had been annotated by eggNOG with the KEGG or GO terms for photosynthesis (KEGG pathway 00195; GO:0015979), with the mean computed for each treatment and species. Pathway visualizations were produced using the pathview package v1.44.0 (Luo & Brouwer, 2013).

### Subgenome-Specific Expression in Hybrids

We used one-to-one orthologs identified by OrthoFinder to compare heat and cold responses between *V. macrocarpon* and *V. oxycoccos* subgenomes within hybrids. To identify one-to-one orthologs, we filtered the OrthoFinder output to retain only gene pairs for which each gene had exactly one ortholog in the other species. DESeq2-normalized counts were subset to include only one-to-one ortholog pairs and separated into *V. macrocarpon* and *V. oxycoccos* subgenomes. These were separated by experiment as described above and analyzed using limma-voom (Smyth, 2005), with results for the interaction term subgenome*treatment used to identify genes with significant (FDR < 0.05) expression divergence between subgenomes under stress. Functional enrichment analyses were conducted on results ranked by t-statistic as described above.

### Expression Differences Among Hybrid Lines

We tested for the effect of genotype (“Line”) using likelihood ratio tests (LRT) within DESeq2, with heat and cold experiments analyzed separately, to identify genes with variable expression among hybrid lines. Pairwise comparisons between each line were also performed to identify genes driving principal component analysis (PCA) separation. An aggregate responsiveness metric was calculated for each line by calculating the mean log fold change for every gene retained by DESeq2 after the filtering methods described above. “Uniquely regulated genes” were defined as genes that were differentially expressed in all other hybrid lines (FDR < 0.05) relative to the focal line. Total ranked outputs were analyzed for functional enrichment by GSEA as described above. Restricted lists of uniquely regulated genes were analyzed by gene ontology enrichment analysis (GOEA) using the enricher() function in clusterProfiler, and enrichments with FDR < 0.05 were considered significant. The background gene universe for GOEA was defined as the set of all genes annotated with a GO term from the concatenated CDS.

### Transgressive Expression in Hybrids

We used the HybridExpress package in R (Almeida-Silva *et al*., 2024) to identify instances of transgressive expression in all four hybrids relative to both parental species. Due to limitations in cultivating *V. oxycoccos* and obtaining sufficient tissue, only ambient temperature conditions were tested, precluding stress treatments. Expression data was filtered to one-to-one orthologs, and expression in hybrids was compared to *in silico* midparent values generated with the add_midparent_expression() function. DESeq2-derived get_deg_list() outputs were categorized into parental, additive, or transgressive modes, and gene lists for each category were evaluated for functional enrichment by GOEA as described above.

## Results

We conducted a time-course RNA sequencing (RNA-seq) experiment using *Vaccinium macrocarpon* (Vmac) and four *V. macrocarpon* x *V. oxycoccos* F1 hybrids to investigate the acute transcriptional basis of temperature responses in cranberry and interspecific hybrids (Figure 1). Plants were exposed to acute heat (40°C and 47°C) or cold (11°C and 3°C) stress following acclimation at 21°C, with RNA collected at three key time points per treatment. A total of 90 RNA extracts (5 plants × 3 biological replicates × 3 timepoints × 2 treatments) were prepared and sequenced using Oxford Nanopore Technologies (ONT) long-read cDNA sequencing. Transcript abundance was quantified by aligning reads to a reference coding sequence database, enabling comparisons across parental and hybrid genotypes. We aimed to identify genes and transcriptional networks responsive to temperature stress, with a particular focus on those showing non-parental expression patterns in hybrids that may underlie temperature resilience traits, by contrasting expression patterns among *V. macrocarpon*, the F1 hybrids, and their inferred parental components.

### Updated parental genomes are highly syntenic

When high-quality genome assemblies are available for both parental species of an F1 hybrid, RNA sequencing (RNA-seq) data can be partitioned and analyzed according to subgenome origin. Recently published reference genomes for *V. macrocarpon* and V*. oxycoccos* (Kawash et al., 2022) provided this opportunity; however, the *V. oxycoccos* genome lacked chromosome-level scaffolding. Given the high structural and gene content similarity between the two species, we scaffolded the *V. oxycoccos* assembly using the chromosome-resolved *V. macrocarpon* genome as a guide to enable ortholog identification. We re-predicted genes for both assemblies using a consistent pipeline to ensure compatibility in downstream RNA-seq analyses. This approach yielded a highly syntenic alignment at the DNA level and revealed conserved orthologs retained from past whole-genome duplication (WGD) events at the protein level (Figure S1). Despite the overall high synteny, approximately 11% (∼3,000) of genes exhibited presence/absence variation (PAV), likely reflecting post-WGD fractionation. Using the chromosome-resolved genomes for read alignment, mean RNA-seq mapping rates were 72.0% for V. oxycoccos, 74.7% for *V. macrocarpon*, and 69.4% for the F1 hybrids.

### Differential expression analysis of *V. macrocarpon* and hybrids considered separately

To establish a baseline understanding of transcriptomic responses to temperature stress, we first evaluated gene-level expression changes within *V. macrocarpon* and hybrid genomes independently. This analysis allowed us to quantify the extent of transcriptional change during stress exposure in each species and to identify enriched functional categories associated with up- or downregulated genes.

We first compared responses to heat stress. Among the hybrids, of 27,730 genes retained for analysis, 3,719 (13.4%) were differentially expressed between time 0 and 30 minutes of heat treatment, including 854 upregulated and 2,865 downregulated genes. 683 gene ontology (GO) terms and 20 Kyoto Encyclopedia of Genes and Genomes (KEGG) pathways were significantly enriched. In *V. macrocarpon*, of 15,392 genes retained, 1,199 (7.8%) were differentially expressed, with 541 upregulated and 658 downregulated. 40 GO terms and 13 KEGG pathways were significantly enriched.

We then compared responses to cold stress. In the hybrid, of 18,681 genes retained for analysis, 175 (0.94%) were differentially expressed, with 143 upregulated and 32 downregulated. 296 GO terms and four KEGG pathways (circadian rhythm, plant hormone signal transduction, photosynthesis, and ribosome) were significantly enriched. In *V. macrocarpon*, of 14,446 genes retained, 270 (1.9%) were differentially expressed, with 201 upregulated and 69 downregulated. 262 GO terms and 10 KEGG pathways, including flavonoid biosynthesis and photosynthesis, were significantly enriched.

### Differential expression analysis of *V. macrocarpon* subgenome

We assessed differential gene expression in the *V. macrocarpon* subgenome in response to heat and cold stress across hybrids and *V. macrocarpon*. This approach isolates the effect of hybridization, enabling the identification of genes and functional categories of the *V. macrocarpon* subgenome that behave differently in a pure versus hybrid genomic background. Throughout, we used GSEA to identify functional differences, as this rank-based method is sensitive to polygenic shifts in gene expression and can detect pathway-level enrichments even when few individual genes are statistically significant.

Only one *V. macrocarpon* gene, *psbA,* which encodes the D1 subunit of photosystem II (PSII), exhibited a significantly divergent response to heat in different genomic backgrounds (Wald test, FDR = 0.00014). *psbA* was upregulated 1.5-fold (Wald test, *p* = 0.026) in hybrids but 17-fold (Wald test, *p* = 3.25 x 10^-25^) in *V. macrocarpon* between 0 and 30 minutes of heat treatment (Figure S2). Gene set enrichment analysis (GSEA) identified 168 differentially enriched GO terms between hybrids and *V. macrocarpon* under heat stress. Enriched terms were associated with fundamental processes such as photosynthesis and ribosome biogenesis, as well as defense response and response to fungus. KEGG pathway enrichment analysis highlighted four differentially expressed pathways: ribosome, photosynthesis, arginine biosynthesis, and biosynthesis of amino acids.

During cold stress, no individual genes were significantly differentially expressed between the hybrids and *V. macrocarpon*. GSEA identified 53 enriched GO terms and 13 enriched KEGG pathways, largely related to photosynthesis and including response to cytokinin. Enriched KEGG pathways included flavonoid biosynthesis, carbon metabolism, ribosome, and photosynthesis.

### Differential Expression and Enrichment Analyses for *V. macrocarpon* vs hybrid Genomes Under Stress

To compare functional responses to temperature stress between *V. macrocarpon* and hybrids, we collapsed gene expression data to KEGG orthology (KO) terms, enabling cross-genome comparisons despite differing gene content. This approach allowed us to assess whether major biological pathways were regulated differently between species and whether these differences diverged from those observed in the *V. macrocarpon* subgenome alone. Several KEGG pathways showed significant differential regulation between hybrids and *V. macrocarpon*, including pathways related to productivity and nutritional content. Only one pathway, Ribosome, was differentially regulated during heat stress (Figure 2b). Photosynthesis displayed differential expression by species under baseline conditions and differential regulation during cold stress (Figure 2a). Eighteen KEGG pathways exhibited differential regulation during cold stress, including photosynthesis (Figure S3), ribosomes, hormone signaling, and flavonoid biosynthesis (Figure 2c). Ribosomes were differentially regulated under both stress conditions. Under heat stress, hybrids broadly up-regulated ribosomal proteins while *V. macrocarpon* marginally suppressed them. During cold stress, both *V. macrocarpon* and the hybrid exhibited suppression of ribosomal proteins, though this downregulation was more pronounced in *V. macrocarpon* (Figure S3). Additionally, flavonoid biosynthesis was minimally impacted during cold stress in hybrids but strongly upregulated in *V. macrocarpo*n. This is of particular interest because the flavonoid biosynthesis pathway is a precursor of anthocyanin biosynthesis, which has a well-characterized role in cold tolerance and is a key determinant of fruit quality (Zalapa 2022; Li & Ahammed, 2023; Kanehisa *et al*., 2025).

**Figure 2:**
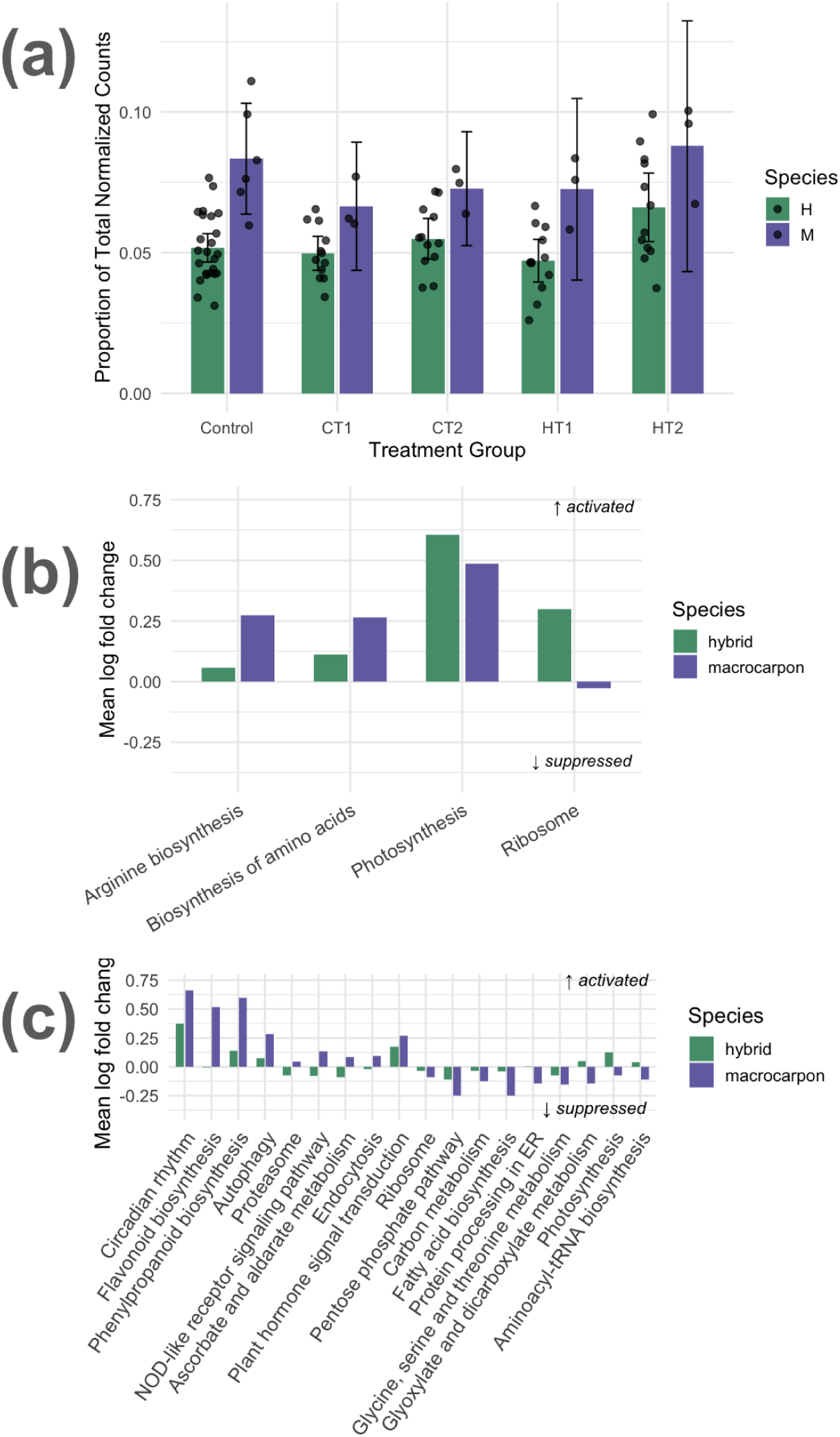
Functional transcriptomic responses to temperature stress in Vaccinium hybrids and V. macrocarpon. A) Total normalized counts for all genes annotated with the pathway or GO term for photosynthesis, by treatment and species (H = hybrid, M = macrocarpon). Error bars represent 95% confidence intervals. B) Expression skew for all KEGG pathways exhibiting significantly different regulation under heat stress, calculated by taking the mean log fold change of all pathway components from 0 to 30 minutes of heat treatment in each species separately. The Ribosome pathway was enriched at FDR < 0.05. The other three enrichments were significant (FDR < 0.05) in the V. macrocarpon subgenome analysis and putatively significant (uncorrected p < 0.05) in the whole-genome analysis. C) Expression skew for all KEGG pathways exhibiting significantly different regulation under cold stress, calculated as in panel B from 0 to 95 minutes of cold treatment. All pathways displayed are significantly differentially enriched by gene set enrichment analysis, FDR < 0.05.

### Differential Expression Analysis of One-to-One Orthologs

To test for subgenome-specific expression patterns, we compared the expression of one-to-one ortholog pairs from *V. macrocarpon* and *V. oxycoccos* within the hybrid genome under control and stress conditions. This allowed us to assess whether either subgenome was constitutively dominant or preferentially activated in response to heat or cold stress. Differential expression analysis of one-to-one orthologs revealed that, under control conditions, 5,339 of 14,184 one-to-one ortholog pairs were significantly differentially expressed; 2,634 of these were *V. oxycoccos*-dominant and 2,705 were *V. macrocarpon-*dominant (Figure 3a). No GO terms or KEGG pathways were identified as significantly enriched by GSEA.

**Figure 3:**
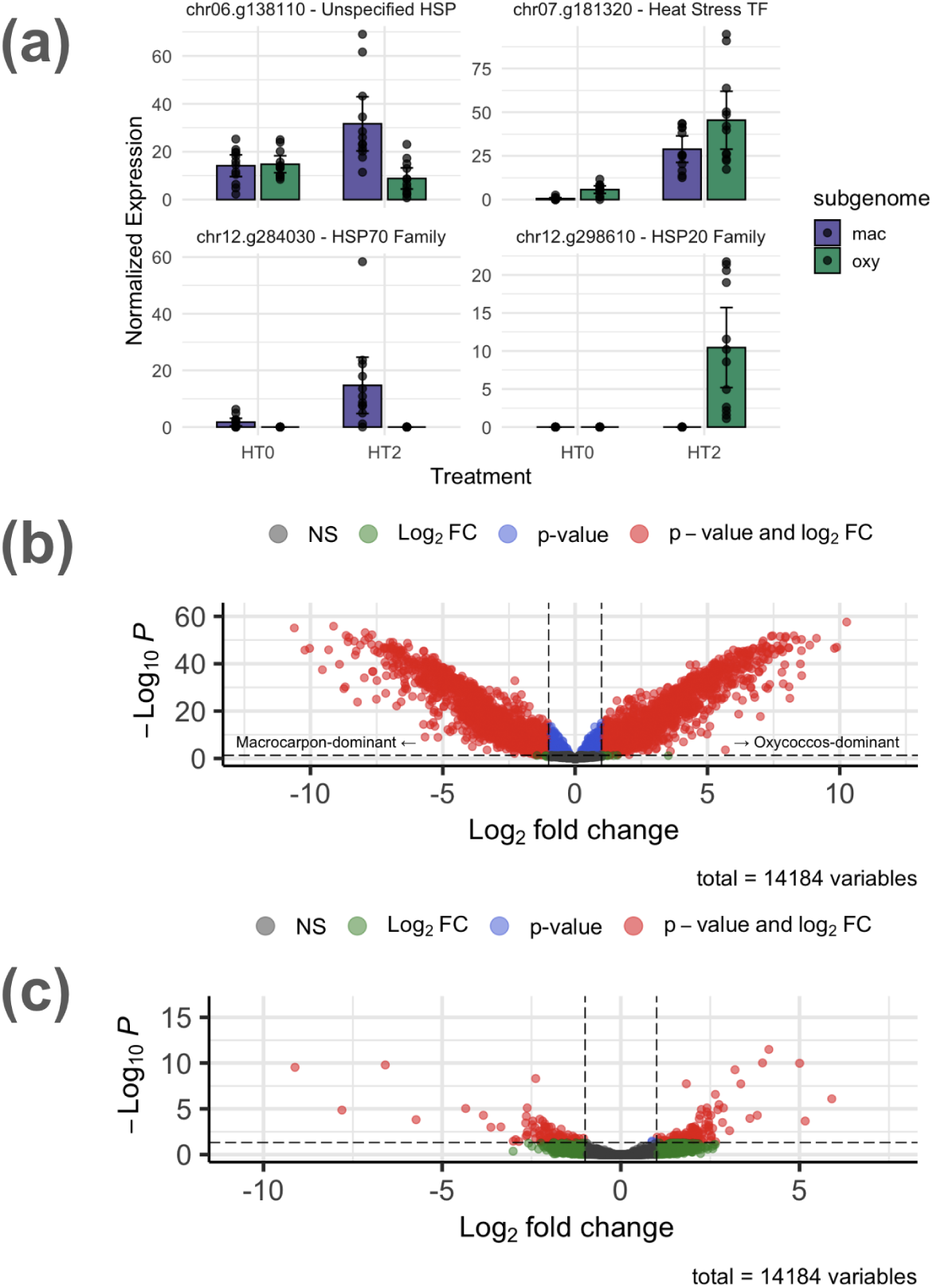
Subgenome-specific regulation of gene expression in Vaccinium hybrids. a) Expression of heat stress genes with significant subgenome effects between control conditions (HT0) and 30 minutes of heat treatment (HT2). b) Volcano plot of the effect of subgenome on expression of all one-to-one ortholog pairs. Negative values indicate V. macrocarpon dominance; positive values indicate V. oxycoccos dominance. c) Volcano plot of the subgenome*treatment interaction term from 0 to 30 minutes of heat treatment, highlighting ortholog pairs that respond differently to heat stress depending on subgenome of origin. Negative values indicate upregulation of V. macrocarpon genes or suppression of V. oxycoccos genes; positive values indicate upregulation of V. oxycoccos genes or suppression of V. macrocarpon genes.

In response to heat, 240 ortholog pairs showed significantly divergent expression changes (Figure 3c). These included three heat shock proteins (HSPs) and one heat shock factor (HSF) (Figure 3a), which showed no pattern favoring either subgenome. However, no enriched GO or KEGG terms were detected by GSEA. Under cold treatment, no ortholog pairs showed expression divergence, and no GO or KEGG terms were enriched.

### Differential Expression and Expression Magnitude Divergence Among Four F1 Lines

We compared expression profiles among four F1 hybrids under heat and cold stress to assess the extent and nature of regulatory diversity introduced through hybridization. Each F1 hybrid line was derived from a different wild *V. oxycoccos* accession crossed to a cultivated *V. macrocarpon* parent; the *V. macrocarpon* parents in two crosses were the cultivar Stevens, and the other two were Pilgrim and Ben Lear (Table 1). By analyzing both the composition and magnitude of expression changes across lines, we aimed to identify variation that could inform future selection, particularly for stress response traits with potential agronomic relevance. Of 27,761 genes retained, 7,717 (28%) showed significant line effects in their response to heat stress, including 76 HSP and HSF. PCA indicated distinct clustering by line and treatment (Figure 4a). PC1, which accounted for 49% of the variance, separated samples by treatment, and PC2, which accounted for 17% of the variance, separated samples by line. GSEA identified 23 enriched GO terms among these genes, related to stress responses (e.g., response to heat and response to oxidative stress) and ribosomes. Four KEGG pathways were significantly enriched: ribosome, protein processing in endoplasmic reticulum, oxidative phosphorylation, and proteasome.

**Figure 4:**
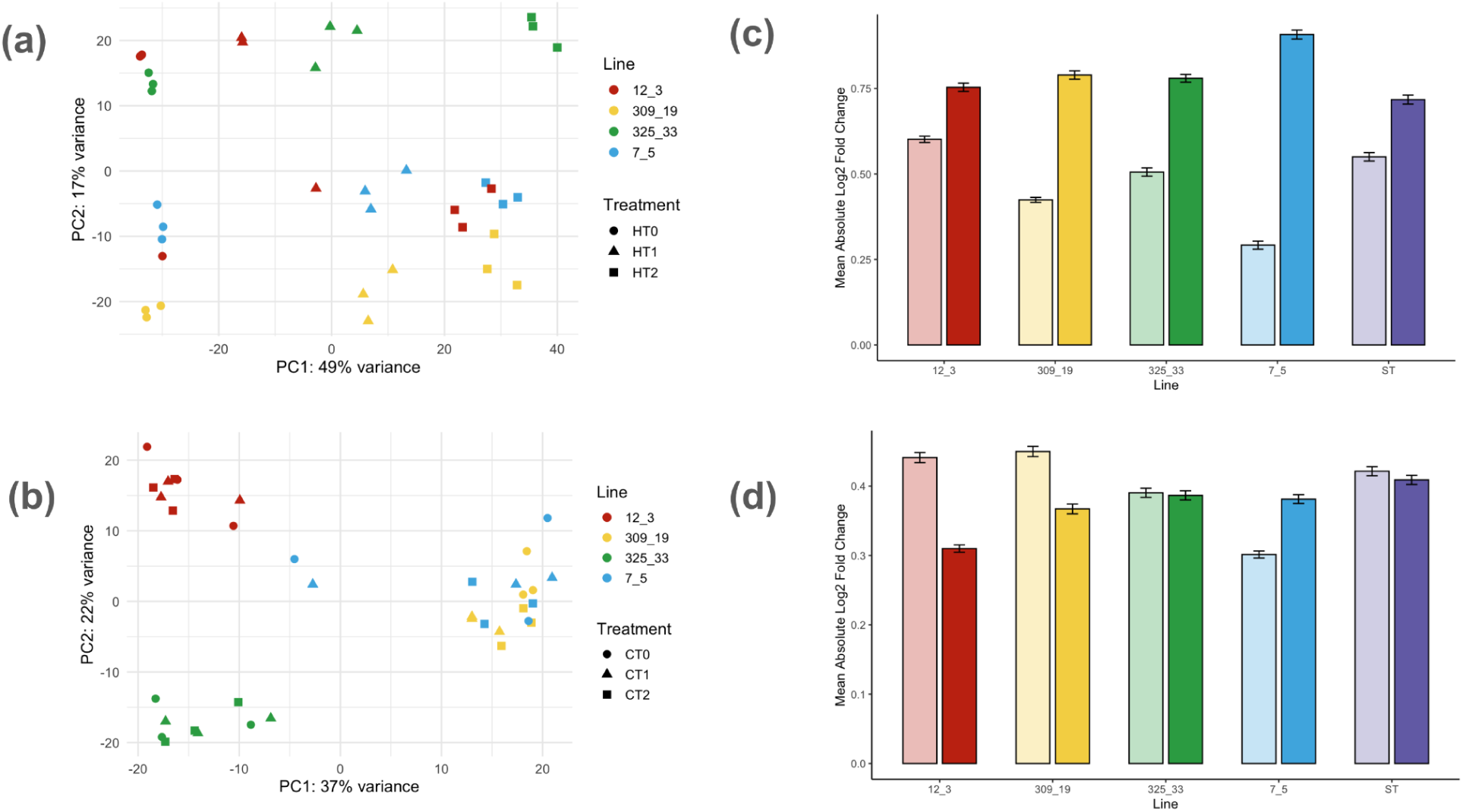
Differences in composition and magnitude of stress responses between hybrid lines. A) PCA of hybrid lines in the heat stress experiment. Note clear separation of treatment conditions along PC1 and fair separation of lines along PC2. B) PCA of hybrid lines in the cold stress experiment. Note distinct clustering of lines but inconsistent separation of treatment conditions. C) Mean absolute log fold change across all genes in DESeq analysis of heat treatment experiment for all hybrid lines and V. macrocarpon (Stevens, ST). Saturated bars represent time 2:time 0 contrast; desaturated bars represent time 1:time 0. Error bars represent 95% confidence intervals. D) Mean absolute log fold change across all genes in DESeq analysis of cold treatment experiment for all hybrid lines and V. macrocarpon (Stevens, ST).

Differential expression between line pairs varied substantially. For example, only 40 genes (0.1%) were differentially expressed between Lines 325_33 and 12_3, which share a Stevens parent, while Line 309_19 and Line 325_33 differed in 2,389 genes (8.6%) (Table S2). Uniquely regulated gene counts also varied: Line 309_19 had 406 uniquely regulated genes, while line 12_3 had only four (Figure S4). GOEA detected five enriched terms in line 309_19’s uniquely regulated genes and 25 enriched terms in line 7_5’s idiosyncratic genes; those in 309_19 lacked an obvious pattern but included negative regulation of cytokine production, while those in line 7_5 were mainly related to mitochondrial function, vitamin biosynthesis, and oxidative stress response.

Lines also differed in their average magnitude of expressional response to heat, measured as the mean absolute log fold change across all genes (Figure 4c). After 30 minutes of heat exposure, *V. macrocarpon* exhibited the smallest magnitude of response (log2FC = 0.72); all hybrid lines exhibited greater responses, with line 7_5 having the largest (log2FC = 0.91). These log fold changes correspond to an average per-gene expression change of 65% for *V. macrocarpon* and 88% for hybrid line 7_5. Interestingly, *V. macrocarpon* had the second-highest magnitude of change after 15 minutes of treatment (log2FC = 0.55) whereas line 7_5 exhibited the weakest response at the same time (log2FC = 0.29).

Under cold stress, of 28,709 genes retained for analysis, 5,265 (18%) showed significant line effects. These included 21 HSP genes, one HSF, a cold shock gene, and three genes encoding cold-regulated 413 plasma membrane proteins. PCA showed modest separation by line, with lines 7_5 and 309_19 overlapping substantially but lines 12_3 and 325_33 separated fully (Figure 4b). Samples did not cluster strongly by treatment group. PC1, which accounted for 37% of the variance, divided samples into two groups by line, with 12_3 and 325_33 separated from 309_19 and 7_5. PC2, which explained 22% of the variance, further separated 12_3 from 325_33. GSEA indicated enrichment of GO terms related to ribosomes and photosynthesis. Only the KEGG pathway “ribosome” was enriched.

As with heat stress, pairwise comparisons revealed variable differentially expressed gene counts. Each comparison revealed between 698 and 865 differentially expressed genes, except 7_5 vs 309_19, which only had nine (Table S2). Uniquely regulated gene patterns were reversed compared to heat stress; 12_3 and 325_33 both had approximately 190 uniquely regulated genes, whereas 309_19 and 7_5 had only two each (Table S4). Uniquely regulated genes in Line 325_33 were enriched for 21 terms, mostly related to vitamin biosynthesis, carbohydrate metabolism, and ribosomes, while uniquely regulated genes in Line 12_3 lacked functional enrichments. Like under heat stress, lines differed in their magnitude of response, though all showed smaller expressional changes than those observed during heat treatment (Figure 4d). Interestingly, across most lines, the magnitude of expressional response was greater at 60 minutes than at 95 minutes of cold treatment, with expressional response in 12_3 and 309_19 declining steeply between 60 and 95 minutes. Interestingly, *V. macrocarpon* exhibited the weakest response to heat treatment at 30 minutes (HT2) but the strongest response to cold treatment at 95 minutes (CT2) relative to all hybrid lines.

### Analysis of Transgressive Expression and Expression-Level Dominance in Hybrids

To evaluate how hybrid gene expression deviates from additive expectations, we examined expression-level dominance and transgressive expression across one-to-one orthologs under ambient conditions. This analysis enables the identification of genes and functional groups for which hybrid expression resembles one parent more than the other or falls outside the parental range, providing insight into transgressive segregation of cryptic molecular traits and highlighting potential sources of hybrid vigor or dysfunction. PCA indicated differentiation between *in silico* hybrids and *in vivo* hybrids (Figure 5a), indicating pervasive nonadditive expression patterns. Expression in *V. oxycoccos* showed substantially higher variability compared to either *V. macrocarpon* or hybrids, and hybrids clustered much more closely with *V. macrocarpon* than with *V. oxycoccos*.

**Figure 5:**
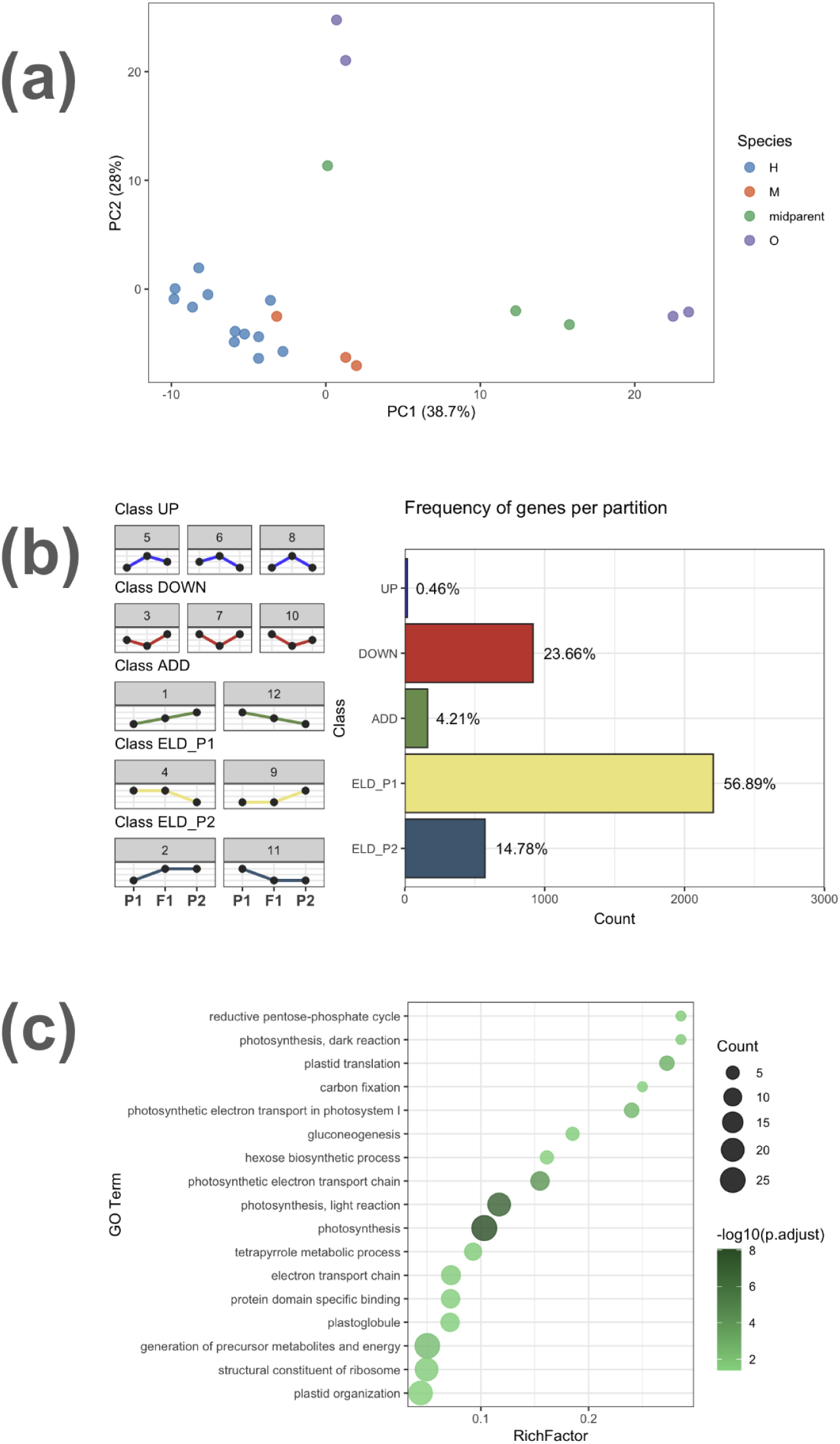
Expression-level dominance and transgressive expression in hybrids relative to parental species. A) PCA of one-to-one ortholog expression for all four hybrids (H), both parental species (M= macrocarpon, O = oxycoccos), and in silico hybrids (midparent; mean expression values between parental species). Note hybrids clustering with V. macrocarpon and strikingly divergent V. oxycoccos expression profiles. B) Frequency of genes per expression-type partition, with potential expression profiles by class at left. Expression-level dominance in favor of V. macrocarpon is the most common expression profile (ELD_P1), with transgressive downregulation (DOWN) and expression-level dominance matching V. oxycoccos (ELD_P2) next. C) GO term enrichments for the DOWN module, indicating potential deficiencies in hybrid function under unstressed conditions. Note depressed expression across several non-overlapping photosynthetic functions.

Most genes showed transgressive expression compared to additive expectations. Expression-level dominance was most common, with 57% of genes matching *V. macrocarpon* expression and 14% matching *V. oxycoccos* (Figure 5b). Genes exhibiting *V. macrocarpon* expression-level dominance were not enriched for any GO terms when analyzed by GOEA. Genes exhibiting *V. oxycoccos* expression-level dominance were enriched for GO terms related to respiration, antioxidant activity, and detoxification. A small proportion (0.51%) of genes exhibited transgressive upregulation, in which expression in the hybrid exceeds that of either parent, but 22.49% showed transgressive downregulation, in which expression in the hybrid is lower than in either parent. Transgressively downregulated genes were enriched for 15 GO terms, most of which were related to photosynthesis (Figure 5c). Transgressively downregulated genes included three HSP genes and one cold acclimation gene (*WCOR413*), potentially indicating deficits in hybrid temperature response.

## Discussion

This study examined the transcriptional responses of *V. macrocarpon* and interspecific *V. macrocarpon x oxycoccos* hybrids to acute heat and cold stress using RNA-seq. We found differences in multiple pathways between *V. macrocarpon* and F1s during stress, both at the whole-organism KEGG ontology level and in the behavior of the *V. macrocarpon* subgenome in a pure versus hybrid genomic context. We found no evidence of coordinated functional subgenomic dominance, but differential expression of orthologous genes in different subgenomes was common. Among F1 hybrids, we observed high variability in expression responses, with many differentially expressed genes and variable magnitudes of average expressional response to stress. Additionally, we observed widespread non-additive expression patterns in hybrids compared to either parental species, with transgressively downregulated genes enriched for photosynthesis-related terms.

### Hybrid responses to temperature stress are characterized by regulatory differences at the pathway level

At the gene level, only one *V. macrocarpon* transcript displayed significant differential regulation between the hybrid and *V. macrocarpon* genomic backgrounds. *psbA,* which encodes the D1 subunit of photosystem II, exhibited modest upregulation in response to heat in hybrids but a sharp increase from low expression in *V. macrocarpon,* reaching similar expression levels after 30 minutes of heat treatment. This difference may be biologically relevant, as overexpression of *psbA* has been demonstrated to improve heat tolerance and photosynthetic performance in *Arabidopsis* (Chen *et al*., 2020). However, whether higher baseline expression in hybrids offers an advantage over the heat-inducible expression observed in *V. macrocarpon* will require further testing.

While few transcripts showed differences individually, gene set enrichment analysis (GSEA) of gene ontology (GO) and Kyoto Encyclopedia of Genes and Genomes (KEGG) terms revealed pronounced divergence between *V. macrocarpon* and hybrid responses, particularly to cold stress. Eighteen KEGG pathways differed significantly between hybrids and *V. macrocarpon* under cold treatment, including fundamental processes such as photosynthesis, and secondary processes like flavonoid biosynthesis and NOD-like receptor signaling. These secondary pathways are especially relevant to cranberry improvement; they underlie anthocyanin production and pathogen defense, respectively, two of the highest priorities for cranberry improvement (Gallardo *et al*., 2018). In contrast, heat stress produced stronger transcriptional responses but fewer functional differences between hybrids and *V. macrocarpon*, primarily involving ribosomal gene expression.

Hybrids appeared to maintain greater metabolic capacity during both types of stress compared to *V. macrocarpon*. Under heat stress, hybrids upregulated ribosomal proteins while *V. macrocarpon* downregulated them. During cold stress, both species suppressed expression of ribosomal proteins, but the degree of downregulation was greater in *V. macrocarpon* than in hybrids. Photosynthesis genes followed a similar pattern during cold stress, being upregulated in hybrids but downregulated in *V. macrocarpon*. These patterns indicate a greater persistence of growth-related functions during stress in hybrids compared to *V. macrocarpon*.

These findings, particularly the metabolic persistence observed in hybrids under stress, support the potential for *V. oxycoccos* germplasm to improve cold tolerance traits in cranberry. The traits implicated here are clearly polygenic, as evidenced by the divergence of entire pathways in the absence of sharp expression changes in most individual genes. Moreover, the pronounced functional differences between hybrids and *V. macrocarpon* during cold stress, compared to the single KEGG enrichment during heat stress, provide more robust evidence for the potential utility of *V. oxycoccos* introgression for improving cold tolerance traits compared to enhancing heat tolerance.

### Subgenomes do not display functional specialization

Subgenome dominance has been observed in F1 hybrids of other plant species, and in some cases has been associated with trait-specific functional specialization (Bird *et al*., 2018; Edger *et al*., 2017). In our data, over a third of orthologous gene pairs displayed expression biases depending on subgenome of origin both during baseline conditions, and 2% exhibited differential regulation under heat stress. However, these biases were not associated with significant functional enrichments of GO terms or KEGG pathways. This suggests that the hybrid genome does not exhibit functional partitioning between subgenomes, but instead reflects allele-specific expression variation distributed across regulatory modules. Such diffuse regulation may lead to functional mismatches if interacting proteins are encoded by divergent parental subgenomes, potentially impairing the efficiency of molecular interactions.

Supporting this interpretation, functional enrichments observed in whole-genome comparisons between *V. macrocarpon* and hybrids closely mirrored those derived from analyses of the *V. macrocarpon* subgenome alone. All KEGG pathways significantly enriched in the *V. macrocarpon* subgenome under cold stress were also enriched in the whole-genome analysis (FDR < 0.05), and all pathways enriched under heat stress were likewise putatively enriched in the whole-genome comparison (uncorrected *p* < 0.05). These results suggest that the *V. macrocarpon* subgenome and the whole hybrid genome behave similarly, with no evidence of compensatory regulation by the *V. oxycoccos* subgenome or partitioning of function between subgenomes.

### Hybrid lines exhibit distinct transcriptional responses during temperature stress

The F1 hybrid lines displayed substantial variation in gene expression responses, indicating that few regulatory features distinct from *V. macrocarpon* are universally shared across hybrids. Nearly a quarter of all genes were differentially expressed in at least one pairwise comparison between hybrid lines, and these variable genes were enriched for agronomically relevant functions. For example, photosynthesis and ribosome-related functions were enriched among variable genes during cold stress, and GO terms related to reactive oxygen species (ROS) response, heat response, and ribosomes were enriched during heat stress. This variation provides a promising source of regulatory diversity that could be refined by selection in breeding programs.

Notably, hybrid lines displayed distinct transcriptional patterns depending on the type of temperature stress. Lines 309_19 and 7_5 each had over 200 uniquely regulated genes during heat stress but only two during cold, while the reverse was observed in lines 325_33 and 12_3. This pattern suggests that a conserved response to one form of temperature stress may be associated with nonstandard regulation under the other. While these findings are based on a small number of hybrid genotypes, they suggest breeding efforts targeting heat and cold tolerance may benefit from stress-specific selection criteria, rather than assuming cross-tolerance to both heat and cold based on general stress response pathways.

Patterns of transcriptional attenuation during stress further support the idea that regulatory restraint may underlie resilience. Under cold stress, *V. macrocarpon* exhibited sustained high-magnitude expression changes, ranking third highest at 60 minutes and first at 95 minutes. In contrast, two hybrid lines showed a spike followed by a decline in expression magnitude, suggesting more efficient acclimation or a less disruptive physiological response. Similarly, after 30 minutes of heat stress, the ostensibly heat-adapted *V. macrocarpon* displayed the smallest transcriptional response of all genotypes compared to the hybrids. These findings suggest that some hybrid genotypes may experience reduced physiological disruption under cold stress but greater sensitivity to heat, further supporting a greater value of hybridization for cold stress tolerance than for heat stress tolerance.

The trade-off between stress defense and productivity is an established challenge in crop improvement (Huot *et al*., 2014, Zhang *et al*., 2020; Ogawa-Ohnishi *et al*., 2022). Excessive suppression of growth and metabolism may impair yield, particularly if recovery is slow, while inadequate defense may lead to damage under acute stress. However, interpreting the adaptive significance of stress-induced expression profiles is difficult. Even among well-characterized stress response genes such as HSPs, the relationship between expression and fitness is complex. Overexpression of HSPs enhances heat tolerance across multiple plant species (Malik *et al*., 1999; Queitsch *et al*., 2000; Park & Hong, 2002). Conversely, heat-adapted genotypes exhibit lower inducibility of HSPs compared to cold-adapted genotypes, potentially indicating a greater reliance on basal mechanisms in heat-adapted plants (Barua *et al*., 2008; Tonsor *et al*., 2008; Zhang *et al*., 2015). Without corresponding physiological data, we cannot determine whether the more transcriptionally responsive hybrids are less well-adapted or more effectively buffered by inducible responses, though the muted expression changes in lower-latitude *V. macrocarpon* compared to the hybrid suggest an adaptive value for attenuated stress responses.

### Hybrids exhibit transgressive downregulation of photosynthetic genes

Under baseline conditions, hybrids showed transgressive downregulation of photosynthetic genes relative to either parent. GO enrichment analysis of transgressively downregulated genes revealed significant depletion across several photosynthesis-related terms, suggesting widespread reductions in photosynthetic activity. During cold stress, photosynthesis genes were upregulated in hybrids but downregulated in *V. macrocarpon*, though total expression was still considerably higher in *V. macrocarpon* than in hybrids across all stress conditions.

The contrasting transcriptional responses to cold may reflect divergent thermal optima for photosynthesis inherited from the parental species. *V. macrocarpon*, native to lower latitudes, may have evolved a higher optimal photosynthetic temperature, leading to photosynthetic repression during cold exposure. In contrast, hybrids with the circumboreal *V. oxycoccos* subgenome may maintain higher performance at lower temperatures. Chilling stress for *V. macrocarpon* may resemble favorable conditions for *V. oxycoccos*, leading to transcriptional divergence.

Alternatively, the low baseline expression of photosynthetic genes in hybrids may reflect cytonuclear incompatibilities (Barnard-Kubow *et al*., 2016). Many photosynthetic and ribosomal proteins are encoded in the nucleus but function in chloroplasts or mitochondria, requiring coordination of nuclear and organelle genomes. Though *V. macrocarpon* and diploid *V. oxycoccos* were reported to have “nearly identical” mitochondrial and plastid genomes (Diaz-Garcia *et al*., 2019), the same study found considerable intraspecific organellar sequence diversity within *V. macrocarpon* and among sympatric *V. oxycoccos*. Thus, mismatches between nuclear and organellar genotypes may still contribute to disrupted expression patterns in hybrids. Supporting this possibility, all four KEGG pathways differentially regulated in the *V. macrocarpon* subgenome between a pure and hybrid background (photosynthesis, ribosome, arginine biosynthesis, and amino acid biosynthesis) involve nuclear–organelle coordination (Winter *et al*., 2015).

Finally, some gene expression differences may result from more general consequences of genomic incongruence. Merging two independently evolved subgenomes can produce unexpected interactions between regulatory elements, disruptions to gene dosage balance, or induced homozygosity of genes lacking orthologs (Birchler & Veitia, 2010). Hybrid weakness has been observed in a variety of intraspecific and interspecific hybrids of important crop plants, including context-dependent weakness during temperature stress (Hermsen, 1967; Chu & Oka, 1972; Shii *et al*., 1980; Chen *et al*., 2014). The transgressive downregulation of stress response genes in this study raises the possibility that similar context-dependent incompatibilities may occur in *Vaccinium* hybrids.

### Implications for crop wild relative (CWR) conservation

Our results underscore the value of *V. oxycoccos* and other CWRs as reservoirs of functional diversity. Genetic variation within *V. oxycoccos* appears to drive substantial regulatory differences between hybrid progeny, with a greater number of differentially expressed genes and enriched functional categories observed among F1 lines than between all hybrids and *V. macrocarpon*. While some differences between F1s may reflect variation among *V. macrocarpon* parents, comparable divergence between two F1s sharing a common Stevens parent (12_3 and 325_33) suggests a strong contribution from *V. oxycoccos* alleles in shaping this diversity.

These findings highlight the need for broad collection and screening of CWR populations to identify germplasm with agronomic potential, particularly for stress response traits that are not easily phenotyped in the field. Because expression of stress-inducible genes is highly environmentally dependent, valuable genetic variants may remain cryptic without rigorous profiling (Alvarez *et al*., 2015; Neyhart *et al*., 2022) and could be lost without proactive conservation measures. Coordinated conservation of CWRs across *in situ* and *ex situ* efforts will be vital for preserving this diversity and enabling its future use for crop improvement (Khoury *et al*., 2020; Engels & Ebert, 2021).

### Limitations and future directions

This study provides an initial characterization of acute temperature stress responses in *V. macrocarpon* and *V. macrocarpon x oxycoccos* hybrids. However, several limitations should guide future work. Most notably, our sampling captures early transcriptional responses, measured 30 minutes after initiating heat treatment and 95 minutes after initiating cold treatment. Because gene expression changes dynamically during stress exposure (Kilian *et al*., 2007), expression profiles under prolonged or repeated stress may differ substantially.

Moreover, transcript abundance is only a partial proxy for cellular physiology. Post-translational modifications, codon usage, and translation efficiency all influence protein levels and function, with the relative contribution of each factor changing during temperature stress (Wu *et al*., 2021; Han *et al*., 2022). In our experiment, the substantial but variable changes in ribosomal gene expression observed under both stress conditions likely impact efficiency of translation. Future integration with proteomic or metabolomic data would provide deeper insight into the functional consequences of these transcriptional shifts.

It is also important to interpret cold and heat responses in light of stress intensity. While both treatments involved substantial and abrupt temperature change, 45°C represents an extreme heat shock with a risk of mortality, whereas 3 °C falls within the natural range experienced by cranberry, even during reproductive development (DeMoranville, 1998; SCEC, 2025). Consequently, observed differences in expression may partly reflect differences in stress severity rather than with differences between cold and heat responses.

Ultimately, the adaptive significance of gene expression changes remains unclear without physiological or fitness data. Both strong and restrained stress responses may confer advantages depending on context, due to an inherent trade-off between growth and stress defense (Huot *et al*., 2014; Zhang *et al*., 2020, Ogawa-Ohnishi *et al*., 2022). Evaluating whether hybrid-specific expression changes truly correlate with stress resilience will be essential for informing cranberry breeding efforts.

### Conclusions

This study provides new insights into temperature stress responses in *V. macrocarpon* and its hybrids with *V. oxycoccos*. Stress-induced expression profiles displayed functional differences in basal pathways including photosynthesis and ribosomes, which reflected greater metabolic persistence during stress in hybrids compared to suppression in *V. macrocarpon*. Cold stress induced smaller differences in the magnitude of expressional change but was associated with greater detectability of functional differences, suggesting that *V. oxycoccos* introgression may be particularly useful for enhancing cold resilience. Though hybrids exhibited differences from *V. macrocarpon* across functional modules, coordinated subgenome dominance was not observed. The substantial variability among F1 lines with different parents highlights the regulatory diversity introduced through hybridization and provides opportunities for selection in further breeding applications. These findings emphasize the importance of conserving *V. oxycoccos* and other CWRs as reservoirs of genetic diversity.

## Acknowledgements

We thank Kristia Adams for expert technical assistance up to and including the RNA isolations, and C. Benjamin Naman and Joseph Kawash for providing comments on the manuscript.

## Author Contributions

JP conceptualized and designed the experiment. JP and EM performed the data collection. AD, NA, and JK designed and performed the data analysis. AD, EM, and JP wrote the manuscript. TPM and JP supervised the project and provided comments on the manuscript.

## Data Availability

Sequence data is available on SRA under accession PRJNA1281617. Transcript quantification files are available on GEO under accession GSE301608. All scripts used in data analysis are available on Github at https://github.com/audickinson/cranberry-2025.

## Supplementary Material

**Figure S1:**
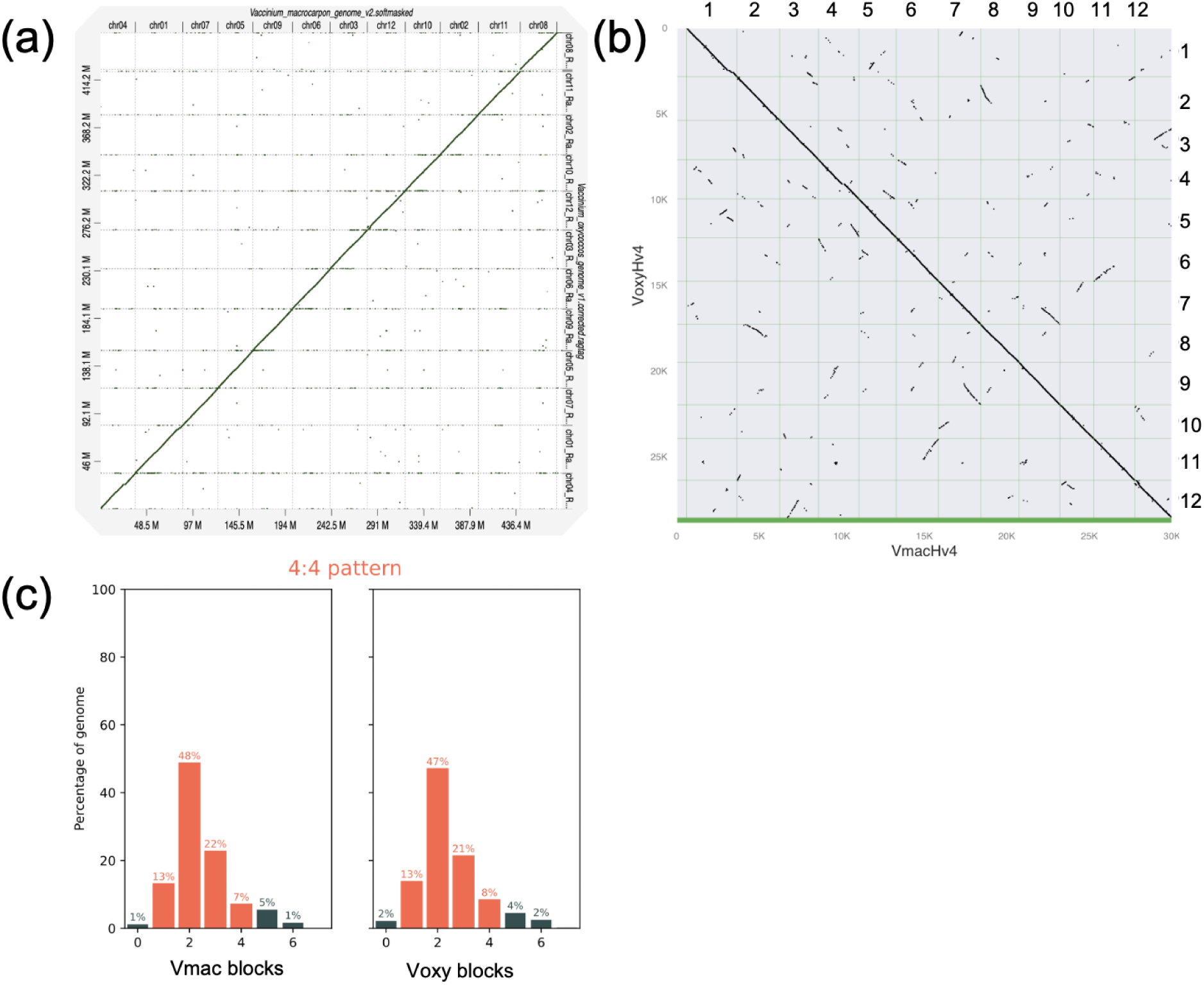
RagTag plot of V. oxycoccos and V. macrocarpon genomes. Dot plot showing syntenic relationship between the V. oxycoccos and V. macrocarpon genome assemblies, generated using RagTag. No major inversions, translocations, or other large-scale structural differences were observed.

**Figure S2:**
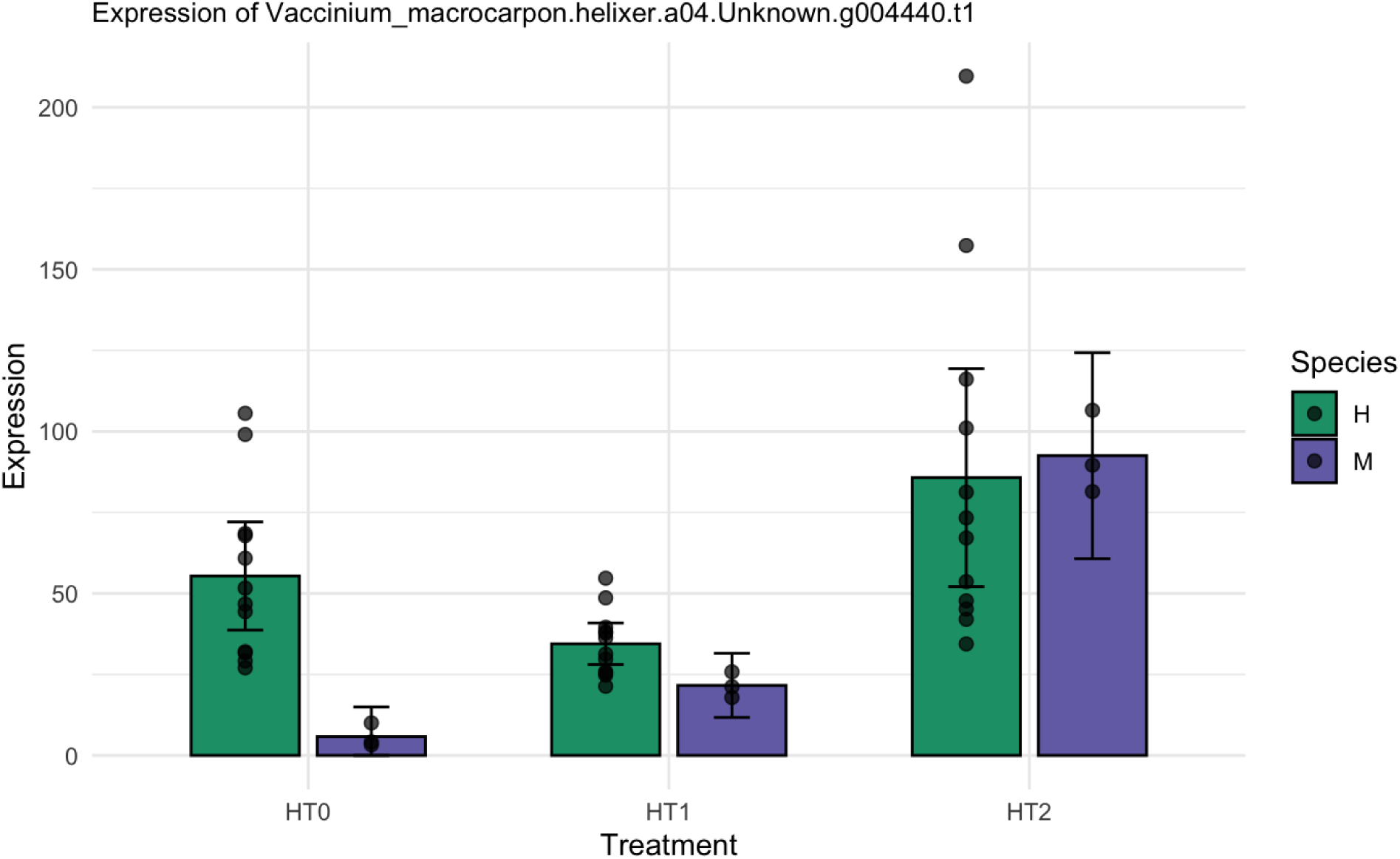
Expression of V. macrocarpon psbA in pure vs hybrid genomic contexts. DESeq2-normalized expression of the psbA gene, which encodes the D1 subunit of photosystem II (PSII), at 0 (HT0), 15 (HT1), and 30 (HT2) minutes after onset of heat treatment in V. macrocarpon (M) and hybrids (H). Error bars represent 95% confidence intervals. While both backgrounds showed upregulation following heat exposure, the response was significantly stronger in V. macrocarpon (17-fold) compared to hybrids (1.5-fold). Additionally, expression declined then increased in hybrids but increased consistently in V. macrocarpon.

**Figure S3:**
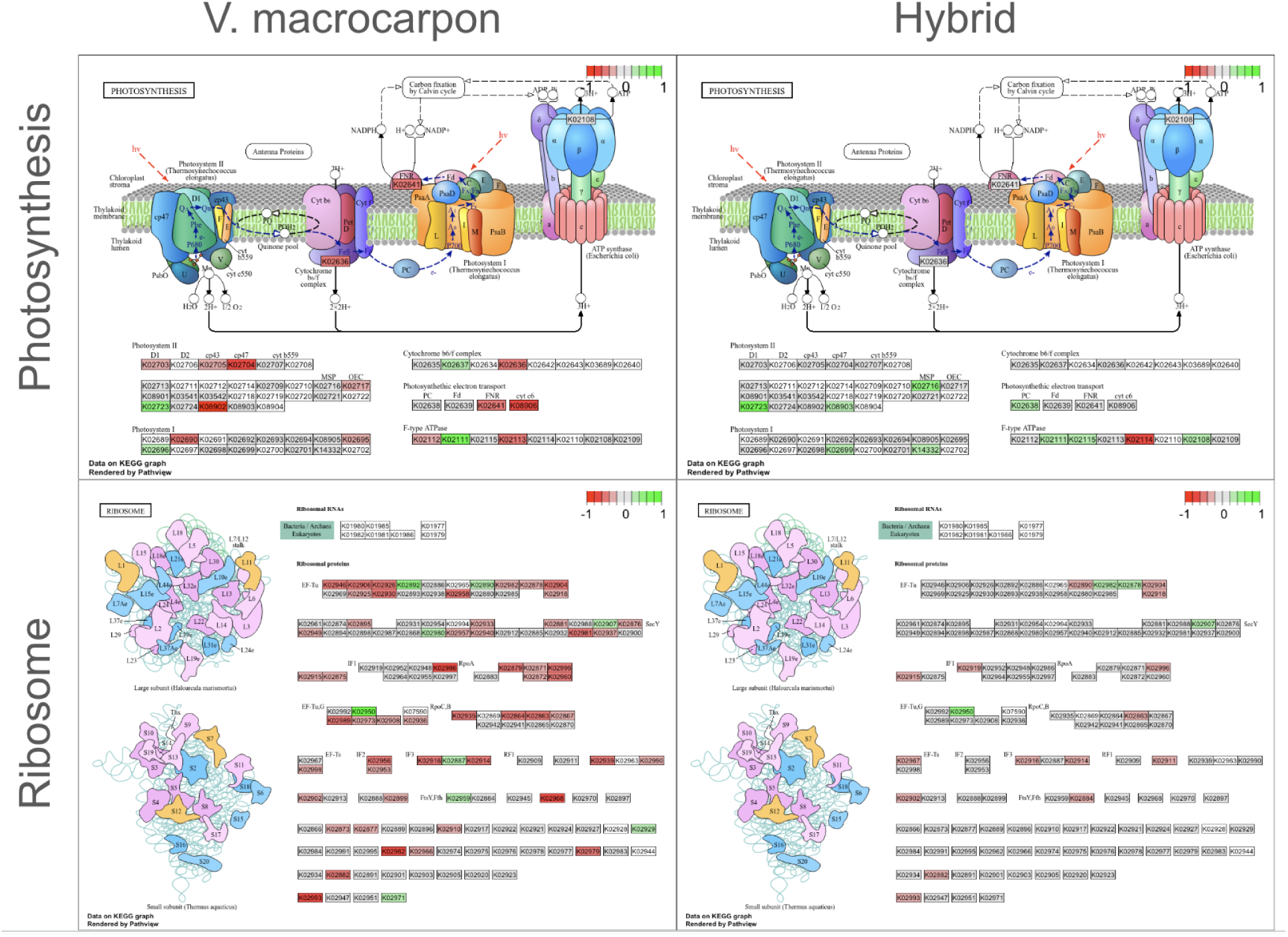
Expression changes of ribosomal and photosynthesis genes in V. macrocarpon and hybrids under cold stress. KEGG pathway diagrams show transcriptional responses of genes annotated to KEGG ribosome and photosynthesis pathways in V. macrocarpon and hybrid genotypes after 95 minutes of cold exposure. Columns represent genotype (pure V. macrocarpon vs hybrid), and rows represent KEGG pathway (Ribosome, Photosynthesis). Colors indicate direction of expression change: red for downregulation, green for upregulation, and grey for no significant change. White indicates no data, either due to lack of expression or lack of annotation. Saturation indicates relative extent of upregulation or downregulation. V. macrocarpon exhibited greater downregulation of both pathways, while hybrids showed more stable or upregulated expression, consistent with enhanced metabolic persistence during cold stress.

**Figure S4:**
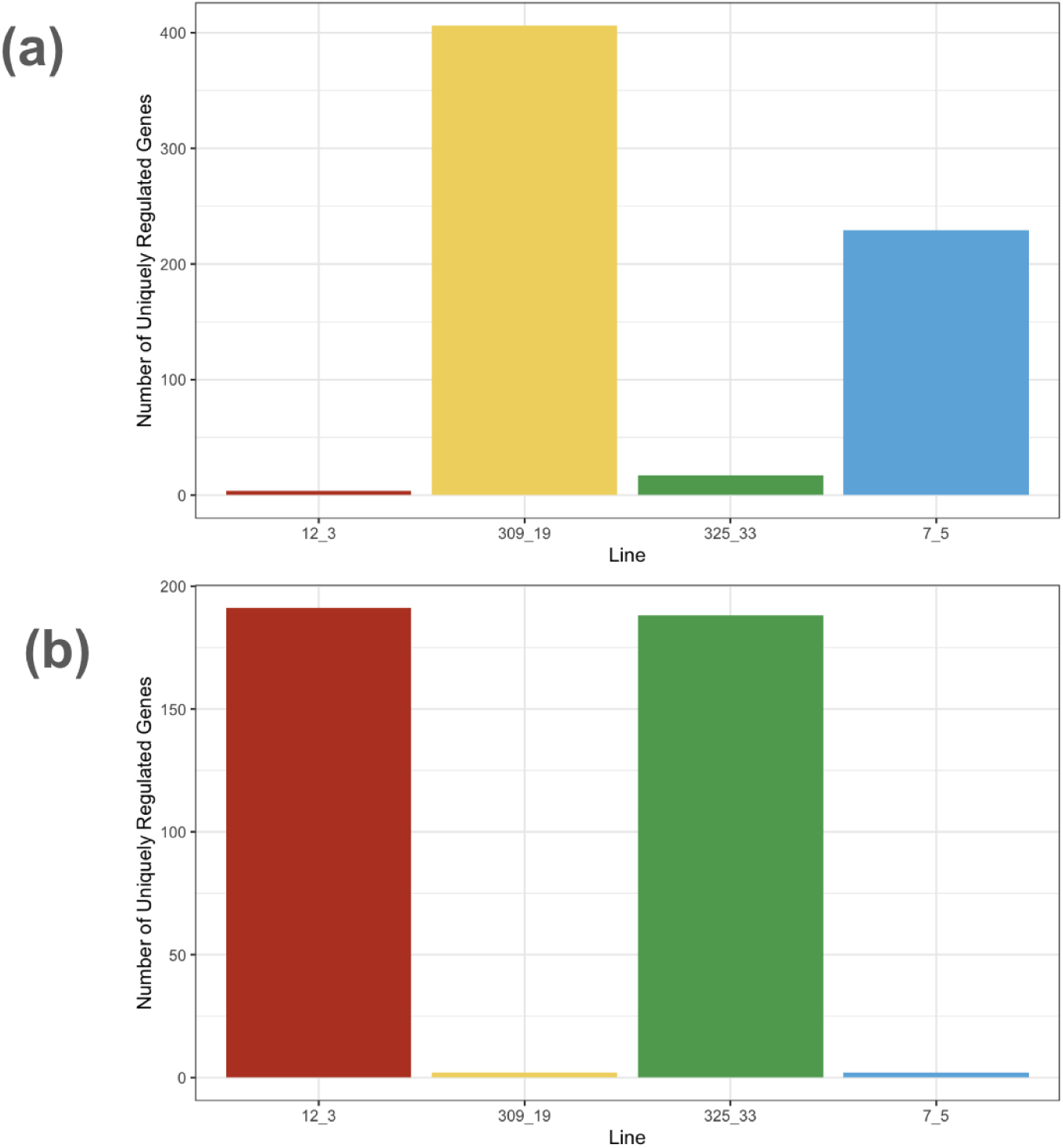
Uniquely regulated genes by hybrid line under heat and cold stress. Bar plots show the number of genes significantly differentially expressed (FDR < 0.05) between the focal line and all other hybrid lines. During heat stress (a), Lines 309_19 and 7_5 exhibited high numbers of uniquely regulated genes, while 12_3 and 325_33 had very few. During cold stress (b), this pattern was reversed, with 12_3 and 325_33 having many uniquely regulated genes and 309_19 and 7_5 having few.

**Table S1:** V. oxycoccos accession information. All accessions were collected in Alaska and belong to populations previously described by Mahy et al. (2000); population codes correspond to those used in that publication. Latitude and longitude indicate the approximate collection site. Accessions listed under “Parents in Hybrid Crosses” were used as parents to create the F1 lines described in Table 1. Accessions listed under “RNA-Seq” were sequenced to enable transgressive expression analysis. These groups differ due to limitations on plant availability.

| Utilization | Plant ID | Collection Location | Population | Latitude | Longitude |
| --- | --- | --- | --- | --- | --- |
| Parents in Hybrid Crosses | NJ96-20 | Between Delta Junction and Glennallen | O2n-1 | 62.2928 | -145.2606 |
|  | NJ96-82 | Cantwell, Alaska | O2n-2 | 63.2313 | -148.5703 |
|  | NJ96-95 | Cantwell, Alaska | O2n-2 | 63.2313 | -148.5703 |
|  | NJ96-127 | Talkeetna, Alaska | O2n-4 | 62.1926 | -150.0634 |
| RNA-Seq | NJ96-21 | Between Delta Junction and Glennallen, Alaska | O2n-1 | 62.2928 | -145.2606 |
|  | NJ96-25 | Between Delta Junction and Glennallen, Alaska | O2n-1 | 62.2928 | -145.2606 |
|  | NJ96-57 | Cantwell, Alaska | O2n-2 | 63.2313 | -148.5703 |
|  | NJ96-112 | Talkeetna, Alaska | O2n-4 | 62.1926 | -150.0634 |

**Table S2:** DE gene counts between hybrid line pairs. Genes were defined as differentially expressed if FDR < 0.05, based on DESeq2 results for the interaction term (treatment*line), with the contrast comparing T2 to T0.

| Line 1 | Line 2 | Number of DE genes (heat) | Number of DE genes (cold) |
| --- | --- | --- | --- |
| 309_19 | 12_3 | 911 | 865 |
| 325_33 | 12_3 | 40 | 816 |
| 7_5 | 12_3 | 921 | 818 |
| 7_5 | 309_19 | 2204 | 9 |
| 325_33 | 309_19 | 2389 | 698 |
| 7_5 | 325_33 | 808 | 749 |

**Table S3:** Differential expression between T2 and T0 for all hybrid lines considered separately and V. macrocarpon. Genes were defined as differentially expressed if FDR < 0.05 and absolute Log2FC >1, based on DESeq2 results for the Treatment term, with the contrast comparing T2 to T0. “All hybrids pooled” refers to the DESeq2 formulation Line + Treatment + Line*Treatment run on expression data from all four hybrid lines. Values for this calculation exceed those in individual line-based contrasts due to increased statistical power.

| Line | Number of upregulated genes, time 0 to 2 |  | Number of downregulated genes, time 0 to 2 |  |
| --- | --- | --- | --- | --- |
|  | Heat | Cold | Heat | Cold |
| 12_3 | 464 | 24 | 302 | 3 |
| 309_19 | 737 | 75 | 717 | 16 |
| 325_33 | 900 | 14 | 1441 | 0 |
| 7_5 | 751 | 89 | 1137 | 15 |
| All hybrids pooled | 854 | 143 | 2865 | 32 |
| ST | 541 | 201 | 658 | 69 |

Supplementary Data S1: Cranberry_RNAseq_ONT_Runs.csv Sequencing run statistics for all sample groups

Supplementary Data S2: Sample_Information.csv Sample information and mapping statistics

